# Systemic Bone Loss Following Myocardial Infarction in Mice is Mitigated by Treatment with a β3 Adrenergic Receptor Antagonist

**DOI:** 10.1101/2020.02.20.958116

**Authors:** Priscilla M. Tjandra, Manali P. Paralkar, Benjamin Osipov, Yi-Je Chen, Fengdong Zhao, Crystal M. Ripplinger, Blaine A. Christiansen

**Affiliations:** Biomedical Engineering Graduate Group, University of California Davis, Davis, CA; Department of Pharmacology, University of California Davis Health, Davis, CA; Department of Orthopaedics, Sir Run Run Shaw Hospital, School of Medicine, Zhejiang University, Hangzhou, China; Department of Orthopaedic Surgery, University of California Davis Health, Sacramento, CA

**Keywords:** Myocardial infarction, Injury/fracture healing, Bone QCT/μCT, DXA, Systems biology - Bone interactors

## Abstract

Myocardial infarction (MI) and osteoporotic fracture (Fx) are leading causes of morbidity and mortality worldwide, and there is epidemiological evidence linking their incidence, suggesting possible crosstalk. MI can exacerbate underlying atherosclerosis through sympathetic nervous system activation and β_3_ adrenoreceptor-mediated release of hematopoietic stem cells (HSCs), leading to monocytosis. We hypothesized that this same pathway may initiate systemic bone loss following MI, since osteoclasts differentiate from monocytes. In this study, MI was performed in 12-week-old male mice (n=24), and mice were randomized to treatment with a β_3_-adrenergic receptor antagonist (SR 59230A) or no treatment for 10 days post-operatively; additional mice (n=21, treated and untreated) served as un-operated controls. Bone mineral density (BMD), bone mineral content (BMC), and body composition were quantified at baseline and 10 days post-MI using DXA; at 10 days post-MI circulating monocyte and neutrophil levels were quantified, and the L5 vertebral body and femur were analyzed with micro-computed tomography. We found that MI led to increases in circulating monocyte and neutrophil levels, but contrary to our initial hypothesis, β_3_-antagonist treatment further increased circulating monocytes compared to untreated mice. BMD and BMC of MI mice decreased at the femur and lumbar spine (−6.9% femur BMD, -3.5% lumbar BMD); β_3_-antagonist treatment diminished this bone loss response (−5.3% femur BMD, -1.2% lumbar BMD). Similarly, trabecular bone volume was decreased in MI mice compared to control mice, and this bone loss was partially attenuated by β_3_-antagonist treatment. These results suggest that MI leads to bone loss at both the axial and appendicular skeleton, and that the sympathetic nervous system may be a modulator of this response. This study is the first to mechanistically show bone loss following an ischemic injury; these findings suggest that bone loss and increased fracture risk may be important clinical co-morbidities following MI or other ischemic injuries.

## Introduction

Osteoporotic fractures (Fx) and myocardial infarction (MI) are two of the leading cause of morbidity and mortality worldwide [1]. Over 20% of patients will die within 1 year following a hip fracture [2, 3], and over 30% will die within 5 years [4, 5]. Similarly, within 5 years of a first MI, 36% of men and 47% of women will die due to MI-related complications [6]. Interestingly, there is strong epidemiological evidence showing that MI is associated with increased risk of subsequent Fx. For example, Gerber *et al*. found that Fx incidence rates increased markedly over time (hazard ratio = 1.32) among those with previous MI compared to control patients [7]. A possible interpretation of these findings is that the incidence of Fx and MI is reflective of advanced stages of underlying chronic diseases such as osteoporosis and atherosclerosis, which are etiologically linked [8]. However, another contributing factor may be that MI initiates an adaptive healing response that actively initiates bone loss systemically, thus increasing subsequent risk of Fx.

Our previous study described a positive feedback loop between Fx and systemic bone loss in mice, which could lead to increased risk of subsequent fractures [9]. In that study, Fx led to significant decreases in whole-body bone mineral density (BMD) in both young (3-month-old) and middle-aged (12-month-old) mice within 2 weeks post-injury. Fx also resulted in ∼11-18% losses of trabecular bone volume in the L5 vertebral body at the same time point. These changes in BMD and bone microstructure were associated with decreased voluntary activity, increased systemic inflammation, and increased osteoclast number and activity at 3 days post-injury. These data demonstrate that acute skeletal injury (femur Fx) initiates a systemic response leading to loss of bone at distant skeletal sites. However, it is not known whether non-musculoskeletal injuries such as MI could also lead to systemic bone loss.

A similar positive feedback loop has been described for the cardiovascular system, wherein the systemic inflammatory response following MI in mice exacerbates underlying atherosclerosis [10]. This study demonstrated that activation of the sympathetic nervous system (SNS) after MI initiated the release of haematopoietic progenitor cells from the bone marrow, ultimately leading to monocytosis, accumulation of monocytes within atherosclerotic lesions, and exacerbated lesion formation. Treatment of mice with a β_3_-adrenergic receptor antagonist after MI lowered protease activity and myeloid cell content, ultimately decreasing the severity of monocytosis. It is possible that a similar SNS-mediated pathway may also be a key mediator of systemic bone loss following Fx or other acute injuries, since osteoclasts have a hematopoetic lineage and differentiate from monocytes. However, the role of the SNS in systemic bone loss following Fx or other injuries has not been described.

In the current study, we sought to determine if acute ischemic injury (MI) leads to a loss of bone systemically, and whether the SNS is a key regulator of this response. We hypothesized that systemic bone loss would occur after MI, and that blockade of β_3_-adrenergic receptors would diminish or prevent this bone loss, implicating the SNS as a mediator of systemic bone adaptation following acute injury. These findings would describe a novel and potentially critical comorbidity associated with MI and other ischemic injuries, and could inform future treatments that aim to preserve skeletal health in these patients.

## Methods

### Animals

Forty-four male C57BL/6 mice were obtained from the Jackson Laboratory (Bar Harbor, ME) at 10 weeks of age and were acclimated to the housing vivarium for 2 weeks prior to the start of experiments. At 12 weeks of age, mice were randomized to MI surgery (n=24) or anesthetized, un-operated controls (n=21). Twenty-one mice (11 MI, 10 control) were administered a selective β_3_-adrenergic receptor (AR) antagonist (SR 59230A, Sigma-Aldrich, St. Louis, MO; twice daily IP injection, 5 mg/kg body weight) for 10 days post-operatively starting immediately post-surgery. All animals were maintained and used in accordance with National Institutes of Health guidelines on the care and use of laboratory animals, and all procedures were approved by the UC Davis Institutional Animal Care and Use Committee.

### Myocardial Infarction Surgery

The left anterior descending (LAD) coronary artery was permanently ligated as previously described [11]. Briefly, mice were anesthetized with isoflurane, intubated, and continuously monitored with a 3-lead electrocardiogram (ECG). A small incision was made, oblique muscles were bluntly separated to expose the ribs, and a small opening was created in the muscle of the 4^th^ intercostal space. The ribs were then separated, and the pericardium was opened. The LAD was identified and permanently ligated using an 8-0 Prolene suture. LAD ligation was confirmed by ST segment elevation on the ECG. The ribs and oblique muscles were closed using a 6-0 Ethilon suture and the skin was closed using wound clips. Approximately 150 μL of sterile saline and 0.1 mg/kg buprenorphine were injected subcutaneously before allowing the mouse to recover in its cage on a 35° C warmer for ∼1 hour. Standard post-operative procedures were followed for 7 days, including analgesia (0.1 mg/kg buprenorphine) twice per day for 48 hours. Wound clips were removed after 7 days. Unoperated control animals were subjected to anesthesia for 30 minutes and followed the same analgesia schedule.

### Measurement of Infarct Size

All mice were euthanized 10 days post-MI, and hearts were removed and placed immediately into cardioplegic solution (composition in mmol/L: NaCl 110, CaCl_2_ 1.2, KCl 16, MgCl_2_ 16, and NaHCO_3_ 10) to prevent continued electrical activity and subsequent ischemic injury to myocytes. Hearts were frozen for 15 minutes, then sliced into 1 mm thick sections (Mouse Heart Slicer Matrix with 1.0 mm coronal section, Zivic Instruments, Pittsburgh, PA). Heart slices were stained with 1% 2,3,5-triphenyltetrazolium (TTC) in PBS for 15 minutes at 35° C after which the slices were stored in PBS for 24 hours. Heart sections were gently blotted with a Kimwipe, then imaged with an office scanner (EPSON Perfection 4990 Photo, Suwa, Japan). Individual color images were taken at 1200 dpi resolution for each heart section, and images were analyzed using ImageJ [12, 13]. To determine the area of ischemic tissue, a color filter was placed on the image to exclude all colors except for white. The filter was then manually adjusted until only the unstained ischemic tissue was highlighted. Total size of ischemic injury was quantified as the total area of ischemic (unstained) tissue in all transverse slices for each heart normalized by the total area of all slices (6-8 sections for each heart).

### White Blood Cell Analysis

Whole blood was collected from the peritoneal cavity at the time of euthanasia for differential white blood cell count to determine the percentage of monocytes and neutrophils in blood. Approximately 0.5 – 0.75 mL of blood was slowly collected through the inferior vena cava using a 30-gauge needle and a 1 mL syringe. The needle tip was removed before the blood was placed directly from the syringe into the collection tube. All blood samples were placed in tubes coated with K_2_EDTA (BD Microtainer^®^, Franklin Lakes, NJ) and were gently inverted ten times before storing at 4° C. Blood samples were transferred to the UC Davis Veterinary Clinical Labs (UC Davis, Veterinary Medical Teaching Hospital) for differential white blood cell count within 24 hours of collection.

### Dual-Energy X-ray Absorptiometry (DXA) Analysis

Whole-body DXA imaging was performed at baseline (one day prior to surgery) and 9 days post-surgery (one day prior to euthanasia) to determine body composition, bone mineral density (BMD) and bone mineral content (BMC) of the whole body, lumbar spine, femoral diaphysis, and whole femur. Mice were anesthetized with isoflurane and placed in a cabinet x-ray system (Mozart^®^, Kubtec Medical Imaging, Stratford, CT). Whole-body analysis automatically excluded the head and wound clips, and BMD, BMC, bone area, lean mass area, and adipose tissue area were calculated using the manufacturer’s software. For the lumbar spine region of interest (ROI), the L4 through L6 vertebrae were manually selected; the whole femur and femoral diaphysis were analyzed using the same method. The femoral diaphysis was determined as the middle third of the femur. The imaging system was calibrated before each use to ensure consistent data.

### Micro-Computed Tomography Analysis

L5 vertebrae and both legs were collected following euthanasia and fixed in 4% paraformaldehyde for 3-4 days before preservation in 70% ethanol. L5 vertebrae and right femora were imaged with micro-computed tomography (SCANCO μCT 35, Brüttisellen, Switzerland) to determine trabecular bone microstructure of the L5 vertebral body and distal femoral metaphysis and cortical bone microstructure of the femoral mid-diaphysis. All bones were imaged according to the guidelines for μCT of rodent bone (energy = 55 kVP, intensity = 114 mA, 6 μm nominal voxel size, integration time = 900ms) [14]. Analysis of trabecular bone in the L5 vertebral body was performed by manually contouring 2D transverse slices in the region between the cranial and caudal growth plates and excluded the vertebral processes. Analysis of the femoral metaphysis was similarly performed with manual contouring beginning at the convergence of the distal femoral growth plate and extending 1500 μm (250 slices) proximal. Trabecular bone volume fraction (BV/TV), trabecular thickness (Tb.Th), trabecular number (Tb.N), and other microstructural parameters were determined using the manufacturer’s analysis software. Analysis of cortical bone in the femoral diaphysis was performed by contouring transverse slices centered on the midpoint of the femur including a total of 600 μm (100 slices). Bone area (B.Ar), cortical thickness (Ct.Th), bone tissue mineral density (TMD) and other microstructural parameters were determined using the manufacturer’s analysis software.

### 3-Point Bending Mechanical Testing of Femora

Mechanical testing was performed on femurs using 3-point bending to determine bone structural and material properties using a materials testing system (ELF 3200, TA Instruments, New Castle, DE, USA). Following μCT imaging, femurs were rehydrated for 10-15 minutes in PBS solution before mechanical testing. The span length of the lower supports was 8 mm, and the femur was positioned so that the posterior aspect of each bone was downward (loaded in tension). The upper loading platen was positioned in the middle of the bone perpendicular to the long axis of the femoral shaft. The bone was preloaded to 1-2 N to ensure contact with the upper platen. Loading was applied at a displacement rate of 0.01 mm/sec until fracture, and displacement and resultant force were recorded at 50 Hz.

Whole-bone structural properties were determined from force-displacement curves using standard methods [15]. Stiffness was calculated as the slope of the linear pre-yield region. Post-yield displacement was determined as the displacement difference between the yield and fracture displacements. Material properties were calculated using previously established beam theory equations [15]. Elastic modulus, yield stress, and ultimate stress were determined using bending moment of inertia (I) and bone radius (c) determined from μCT analysis of the femoral mid-diaphysis.

### Statistical Analysis

All results are expressed as mean ± standard deviation. Cross-sectional data were analyzed by two-way analysis of variance (ANOVA) stratified by operation (MI or Control) and treatment (β_3_-AR antagonist or untreated) to determine main effects and interactions. DXA data were longitudinal and were analyzed using repeated measures ANOVA to determine differences in the time course of outcomes. Post hoc analyses were performed using Tukey’s Honest Significant Difference test. Two-way ANOVA was performed using JMP Pro 14.2.0 (SAS Institute Inc., Cary, NC, USA). Repeated measures ANOVA values were through jamovi (version 0.9). Statistically significant differences were identified at p ≤ 0.05; trends were noted at p ≤ 0.10.

## Results

### Measurement of Infarct Size

Presence of ischemic (unstained) tissue was consistently observed in the left ventricle and areas inferior to the ligation site, confirming successful MI (Fig. 1b). When infarct areas were normalized by total heart area (IA/TA), there was no significant difference between the infarct sizes of the β_3_-AR antagonist treated mice and that of untreated mice (Fig. 1a).

**Figure 1:**
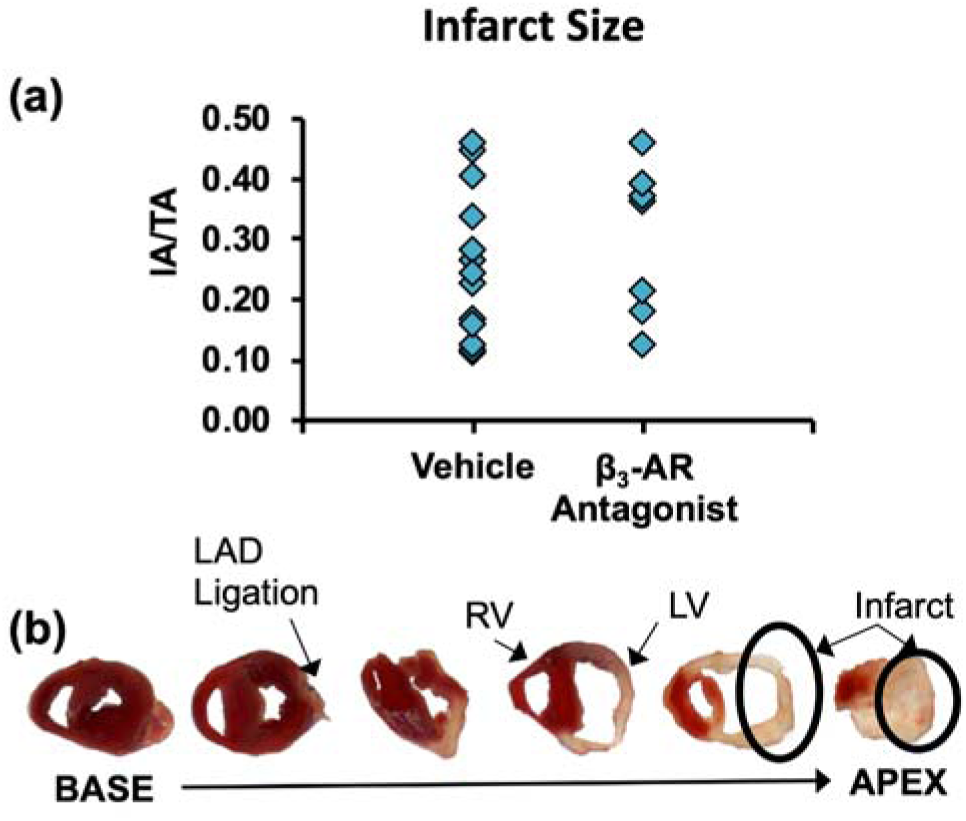
**(a)** Infarct sizes normalized by total heart area for mice following surgical creation of MI. No differences were observed between untreated mice and mice treated with a β3 adrenergic receptor antagonist (IA/TA = Infarct Area/Total Area). **(b)** Representative 1 mm thick cross-sections of a mouse heart 10 days after MI surgery.

### White Blood Cell Analysis

At 10 days post-MI, circulating monocyte levels were significantly greater in MI mice than in Control mice (Fig. 2a; main effect of MI: p = 0.007), and mice treated with the β_3_-AR antagonist had a higher percentage of monocytes in the serum than untreated mice (main effect of treatment: p = 0.033). Similarly, neutrophil levels in MI mice were higher than in Control mice (Fig. 2b; main effect of MI: p = 0.0003), though there was no significant effect of treatment on neutrophil levels. No significant interaction was observed between MI and β_3_-AR antagonist treatment for either monocyte or neutrophil levels.

**Figure 2:**
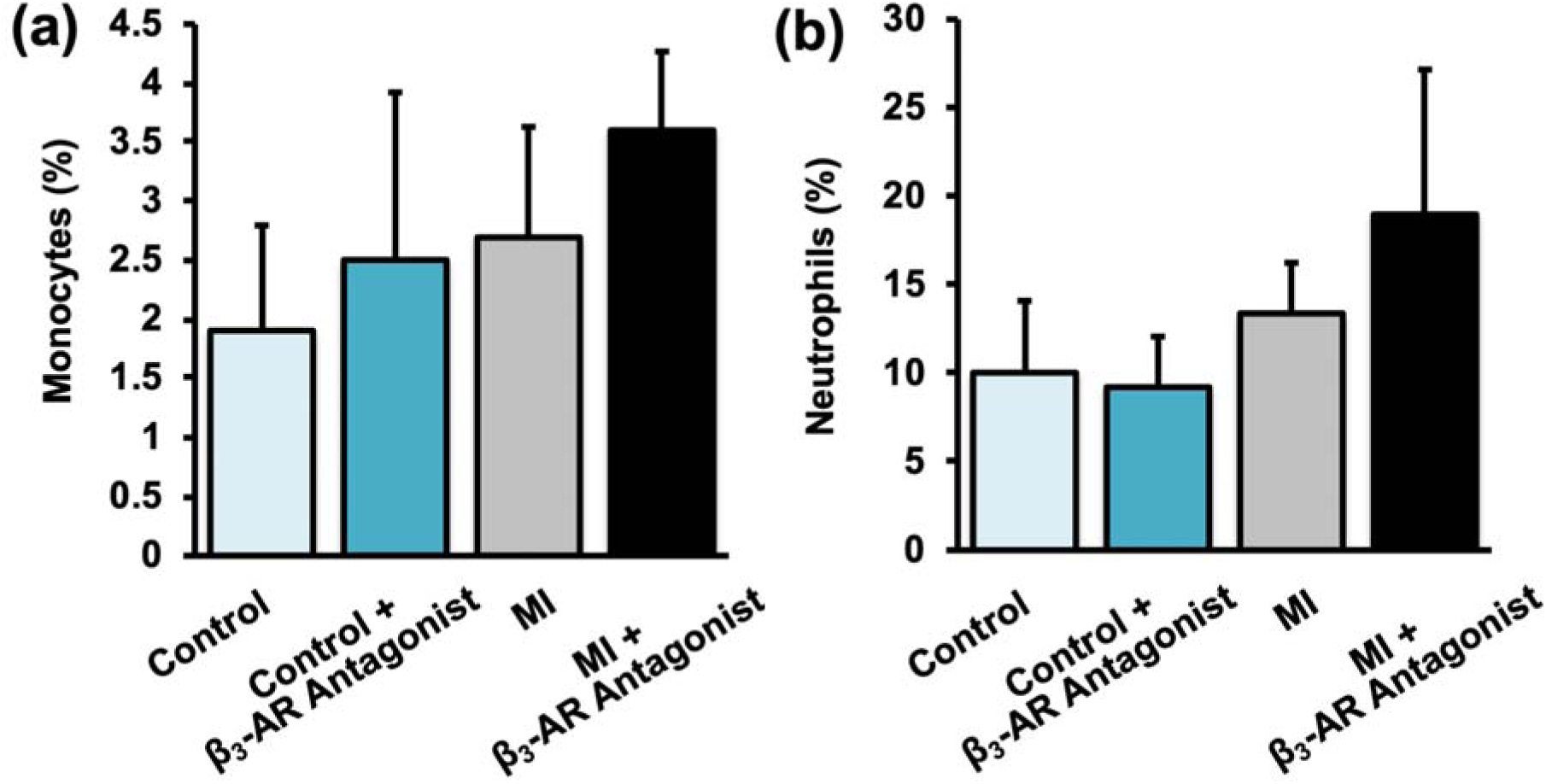
Percent of **(a)** monocytes and **(b)** neutrophils in blood 10 days post-MI. Monocyte levels were significantly increased by MI (main effect of MI: p = 0.007), and by treatment with a β_3_-AR antagonist (main effect of treatment: p = 0.033). Similarly, neutrophil levels were significantly increased by MI (main effect of MI: p = 0.0003).

### Dual-Energy X-ray Absorptiometry (DXA) Analysis

Whole-body DXA of mice revealed few significant changes from baseline to 9 days post-MI in any of the experimental groups, and no statistically significant differences based on MI or treatments (Fig. 3). Generally, whole-body BMD decreased in MI mice from baseline to 9 days post-MI (though not statistically significant), while BMD of Control mice increased significantly during this time period. Whole-body BMC followed similar trends, with MI mice exhibiting less of an increase from baseline than Control mice on average.

**Figure 3:**
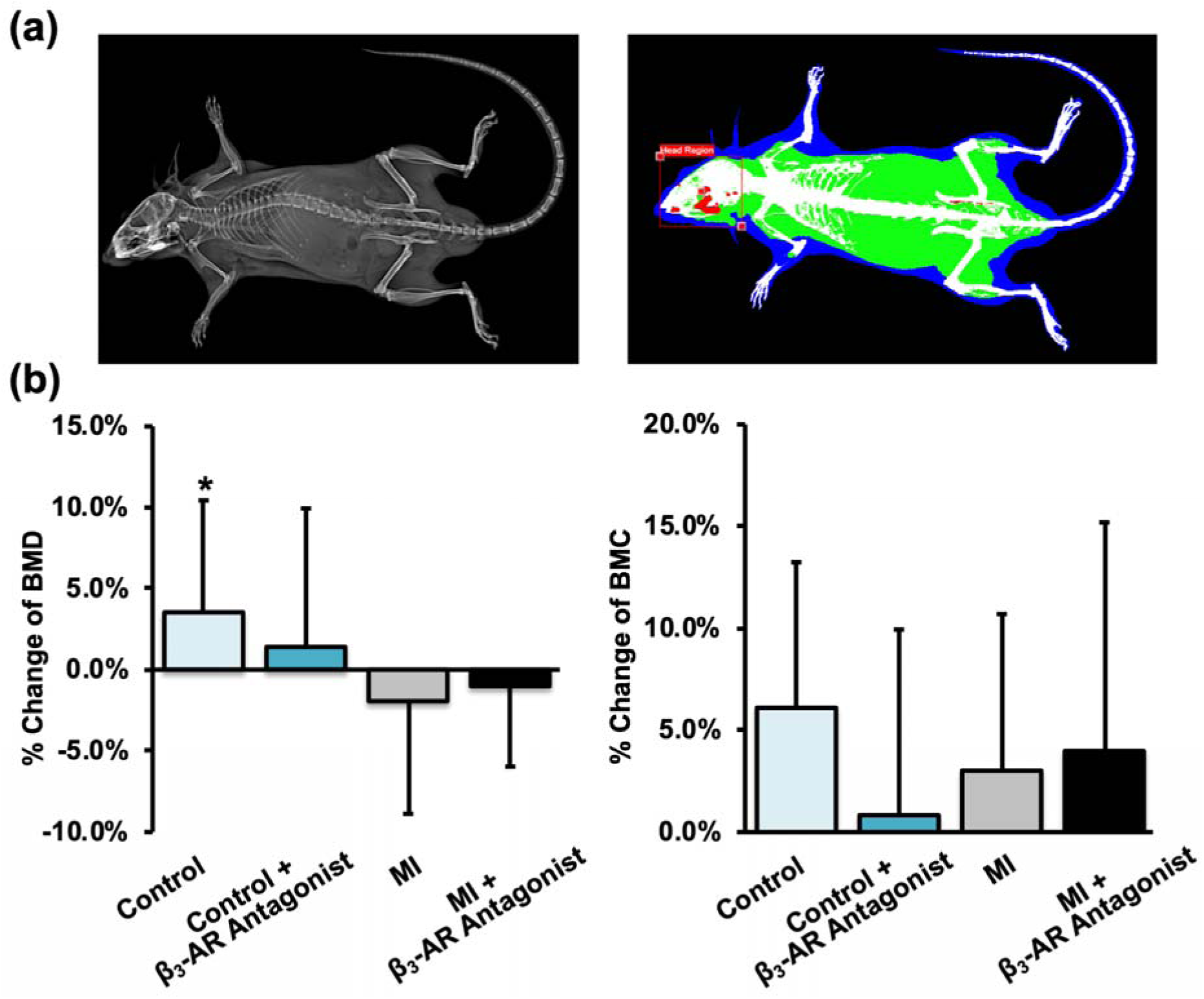
**(a)** X-ray *(left)* and DEXA (*right)* images of the whole mouse. **(b)** Average change of whole-body BMD *(left)* and BMC *(right)* from baseline to 9 days post-MI. * denotes significant change (p ≤ 0.05) from baseline to 9 days post-MI.

Results from analysis of BMD and BMC in the lumbar spine were more definitive (Fig. 4). At this skeletal site, Control mice generally showed an increase in BMD and BMC, while MI mice showed decreases in BMD and BMC from baseline to 9 days post-MI. Additionally, untreated mice exhibited greater changes from baseline (+2.5% BMD and +6.0% BMC for Control mice, - 3.5% BMD and -4.2% BMC for MI mice) compared to β_3_-AR antagonist treated mice (+0.4% BMD and +3.7% BMC for Control mice, -1.2% BMD and -2.5% BMC for MI mice).

**Figure 4:**
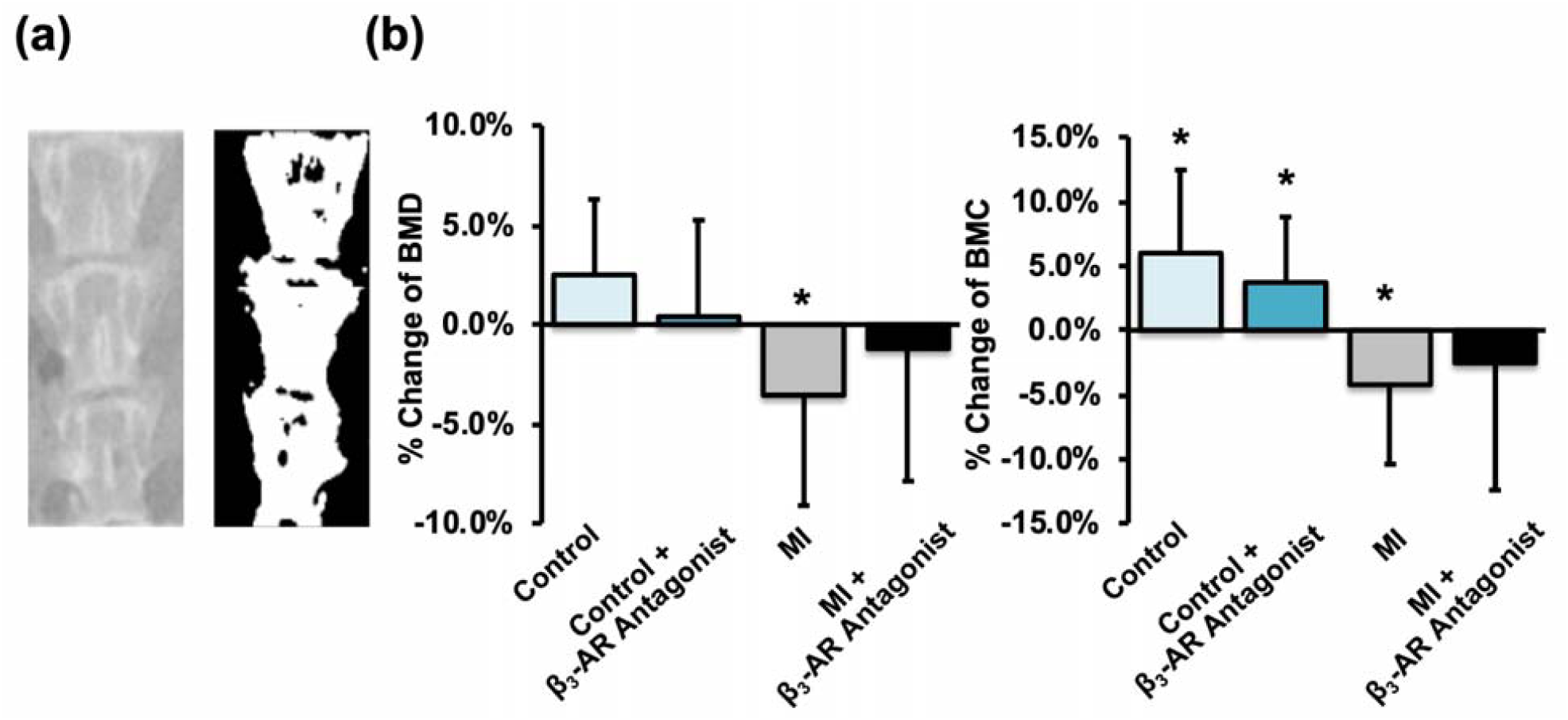
**(a)** X-ray (*left*) and thresholded (*right*) images of the lumbar spine (L4-L6). **(b)** Average change of lumbar BMD (*left*) and BMC (*right*) from baseline to 9 days post-MI. * denotes significant change (p ≤ 0.05) from baseline to 9 days post-MI. Overall, average lumbar BMD and BMC of MI mice decreased from baseline to 9 days post-MI, and untreated mice exhibited greater changes than β_3_-AR antagonist treated mice.

Results from analysis of BMD and BMC of the femur followed similar trends to those from the lumbar spine. For both the whole femur and femoral diaphysis (Fig. 5a and 5c), BMD in the MI groups decreased significantly and this decrease was greater for untreated mice (−6.9% whole femur, p = 0.017; -5.6% diaphysis, p = 0.040) than for β_3_-AR antagonist treated mice (−5.3% whole femur, p = 0.034; -4.5% diaphysis, p = 0.081). No significant differences were observed between β_3_-AR antagonist treated Control mice and untreated Control mice. BMC of the whole femur showed similar results, with BMC of untreated MI mice decreasing from baseline to 9 days post-MI (−5.2% whole, p = 0.018; -6.5% diaphysis, p = 0.012); this change in BMC was mitigated in β_3_-AR antagonist treated MI mice.

**Figure 5:**
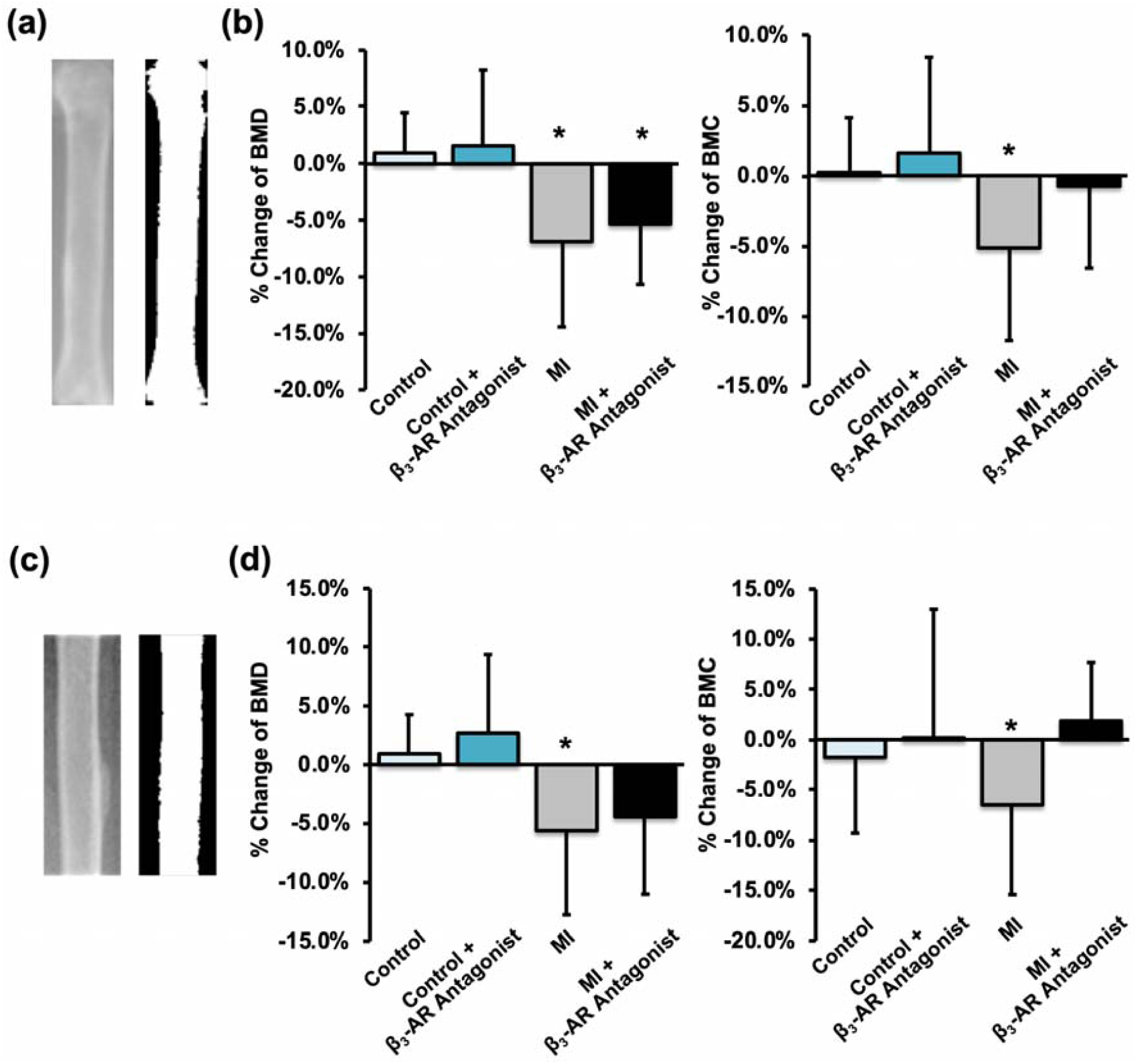
**(a**,**c)** X-ray and thresholded images of the whole femur (a) and the femoral diaphysis (c). (**b**,**d**) Average change of femoral BMD (*left*) and BMC (*right*) from baseline to 9 days post-MI for the whole femur and femoral diaphysis. * denotes p ≤ 0.05 between baseline and 9 days post-MI. Both BMD and BMC decreased significantly in MI mice, and this decrease was greater for untreated mice than for β_3_-AR antagonist treated mice.

Whole-body adipose tissue area decreased from baseline to 9 days for all groups (Fig. 6a), though this change was not statistically significant for untreated Control mice. Whole-body lean tissue area, however, decreased only in MI mice (main effect of MI: p < 0.001), and this change was greatest for β_3_-AR antagonist treated MI mice (−18.3%, p = <0.001).

**Figure 6:**
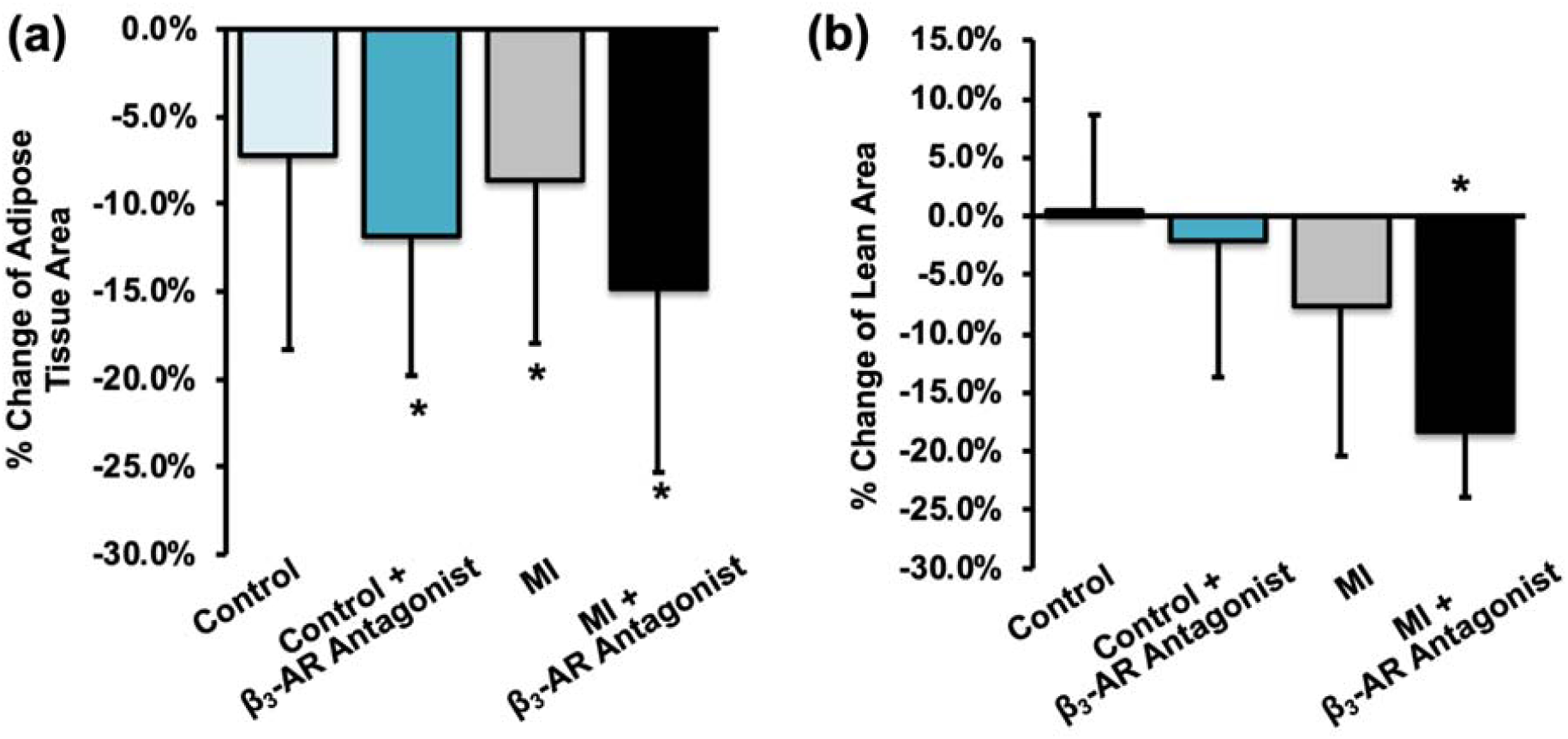
Average change in **(a)** whole-body fat area and **(b)** whole-body lean tissue area between baseline and 9 days post-MI. * denotes p ≤ 0.05 between baseline and 9 days post-MI. Fat area decreased all groups, but lean tissue area decreased only in MI mice (main effect of MI: p < 0.001), and this change was greatest for β_3_-AR antagonist treated MI mice (−18.3%, p = <0.001).

### Micro-Computed Tomography Analysis

Trabecular bone analysis of the axial (L5 vertebral body) and appendicular (distal femoral metaphysis) skeleton yielded results that were generally consistent with those from DXA analysis (Fig. 7). At the L5 vertebral body we observed several statistically significant main effects of β_3_-AR antagonist treatment, with treated mice exhibiting decreased BV/TV (p < 0.001), Tb.N (p = 0.003), Tb.Th (p = 0.045), and apparent BMD (p < 0.001), and increased Tb.Sp (p = 0.001) relative to untreated mice. At the distal femoral metaphysis we observed a significant main effect of β_3_-AR antagonist treatment on Tb.N (p = 0.028) only, with treated mice exhibiting decreased Tb.N relative to untreated mice.

**Figure 7:**
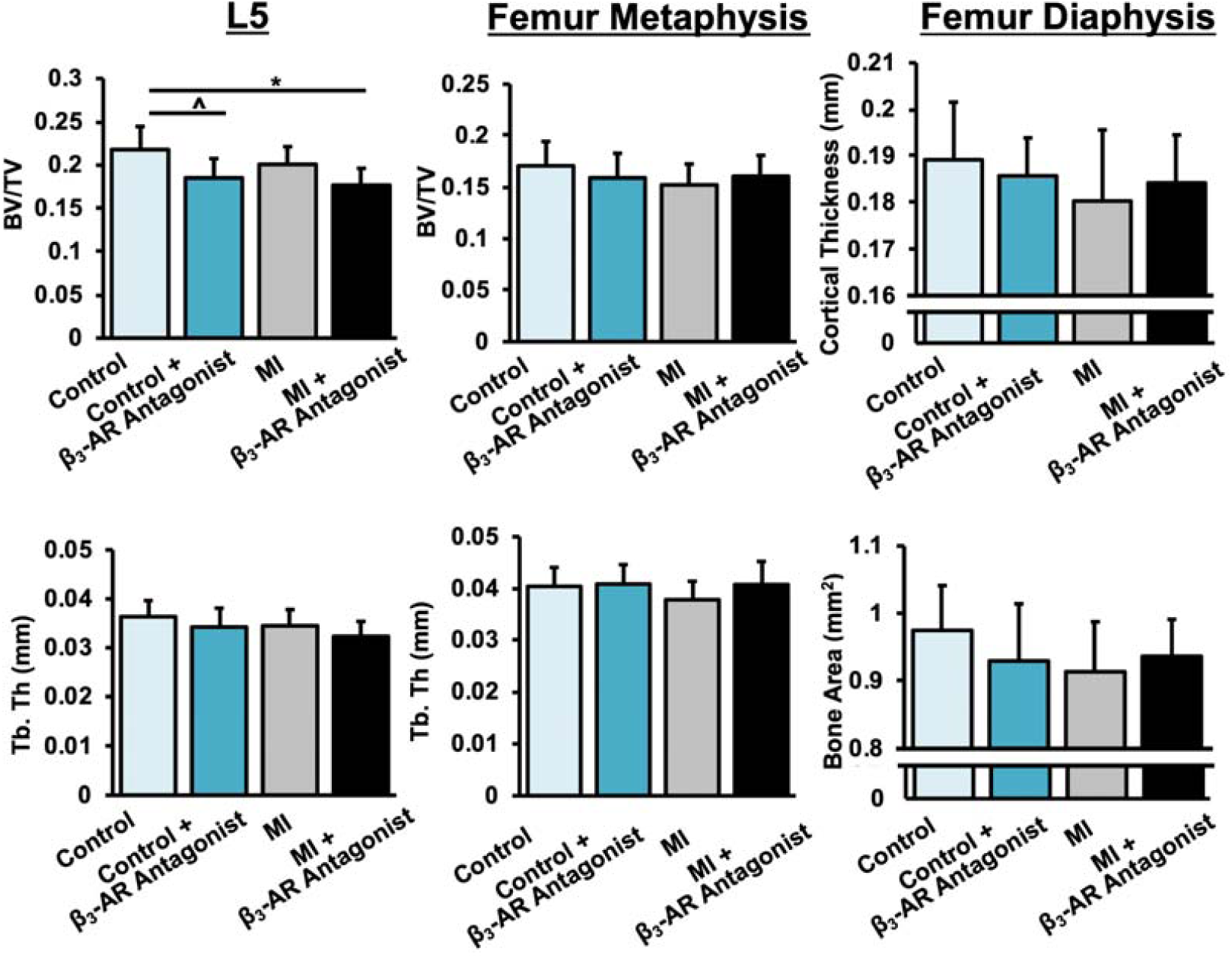
Trabecular bone volume fraction and trabecular thickness of trabecular bone in the L5 vertebral body (left) and distal femoral metaphysis (center); cortical thickness and cortical bone area of the femoral mid-diaphysis (right) 10 days post-MI. BV/TV and Tb.Th were lower in MI mice, and these MI-associated differences were largely mitigated in β_3_-AR antagonist treated mice. p-values < 0.05 are marked by lines between groups.

Untreated MI mice exhibited an 8.2% lower BV/TV in the L5 vertebral body than untreated Control mice (p = 0.103); untreated MI mice also exhibited an 11.4% lower BV/TV (p = 0.051) and a 6.6% lower Tb.Th (p = 0.094) in the distal femoral metaphysis than untreated Control mice. These MI-associated differences were largely mitigated in β_3_-AR antagonist treated mice, though this may be in part due to β_3_-AR antagonist treated Control mice having lower BV/TV and Tb.Th than untreated Control mice. No significant differences were observed between any experimental groups for cortical bone microstructural outcomes at the femoral diaphysis.

### 3-Point Bending Mechanical Testing of Femora

Mechanical testing of femurs in 3-point bending revealed no significant differences between groups in tissue material properties such as modulus of elasticity and ultimate stress (Suppl. Table 1). Similarly, we observed no significant differences in structural properties such as stiffness, ultimate force, and post-yield displacement. However, we observed significant interactions between MI and β_3_-AR antagonist treatment for yield force (p = 0.004) and yield stress (p = 0.046).

**Supplemental Table 1:** Results from 3-point bending mechanical testing of femurs.

|  | **Control** | **Control +**<br>**$\beta_3$-AR Antagonist** | **MI** | **MI +**<br>**$\beta_3$-AR Antagonist** |
| --- | --- | --- | --- | --- |
| **Stiffness (N/mm)** | 126.9 ± 20.2 | 126.8 ± 9.0 | 123.0 ± 13.4 | 115.0 ± 26.6 |
| **\* Yield Force (N)** | 14.0 ± 2.2 | 11.3 ± 1.6 | 12.1 ± 1.6 | 12.6 ± 1.4 |
| **Ultimate Force (N)** | 18.4 ± 2.5 | 17.2 ± 1.8 | 17.9 ± 1.9 | 16.8 ± 1.8 |
| **Post-Yield Displacement (mm)** | 0.17 ± 0.09 | 0.17 ± 0.10 | 0.24 ± 0.18 | 0.13 ± 0.07 |
| **Modulus of Elasticity (kPa)** | 7789 ± 1491 | 8573 ± 1144 | 8344 ± 1258 | 7841 ± 2154 |
| **\* Yield Stress (kPa)** | 112.9 ± 14.9 | 97.2 ± 13.6 | 103.7 ± 19.6 | 108.7 ± 14.1 |
| **Ultimate Stress (kPa)** | 148.5 ± 15.3 | 148.0 ± 8.2 | 153.8 ± 22.1 | 143.9 ± 17.2 |

\* Significant interaction between MI and $\beta_3$ -AR Antagonist treatment ( $p < 0.05$ )

### Correlations of Bone Outcomes with Infarct Size

No significant correlations were observed between infarct size and monocyte or neutrophil levels, or between infarct size and DXA data for whole-body or regional measurements. However, we observed significant negative correlation between infarct size and L5 BV/TV of untreated MI mice and femoral metaphysis BV/TV of β_3_-AR antagonist treated MI mice (Suppl. Fig. 1).

**Figure S1:**
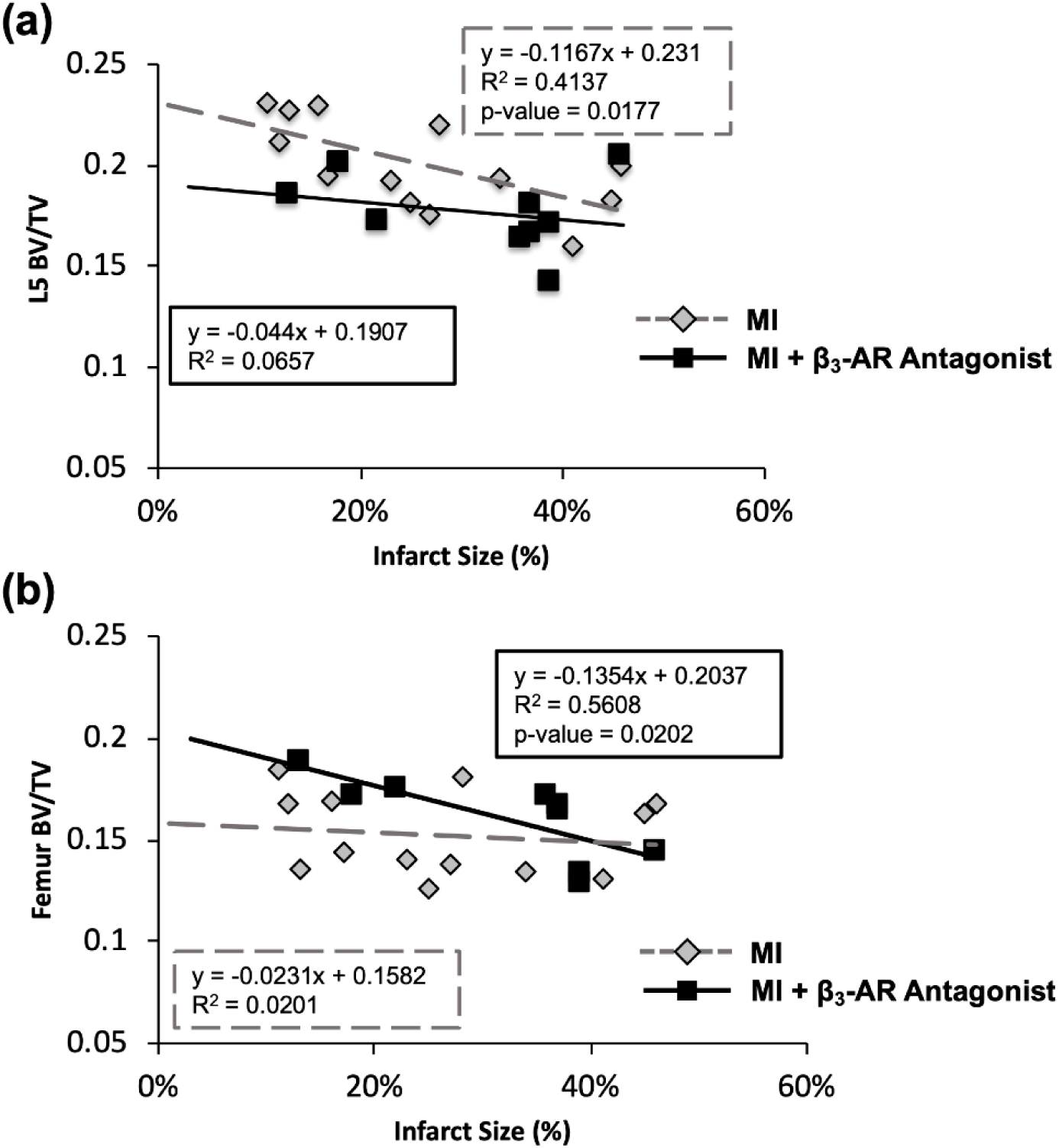
Correlations between ischemic tissue area and (a) L5 BV/TV and (b) femur metaphysis BV/TV. Infarct size was significantly negatively correlated with BV/TV of untreated MI mice at the L5 vertebral body and BV/TV of β_3_-AR antagonist treated MI mice at the distal femoral metaphysis.

## Discussion

In this study, we investigated bone loss in the whole-body and at axial and appendicular skeletal sites following MI in mice. Consistent with our hypothesis, MI mice had a lower BMD and BMC in the axial and appendicular skeletal sites and lower BV/TV and Tb.Th at the same skeletal sites compared to Control mice. BMD and BMC changes in β_3_-AR antagonist treated mice showed a muted effect compared to untreated mice. These results are the first to show a causative effect of MI leading to systemic bone loss, and these data suggest that the sympathetic nervous system may be an important regulator of this response.

The magnitude of systemic bone loss we observed in this study following MI is consistent with our previous study of bone loss following femoral fracture in mice [9]. In both cases, whole body BMD and BMC decreased in injured mice compared to age-matched control mice within 2 weeks post-injury. Additionally, trabecular bone microstructure was diminished in the L5 vertebral body and distal femur in both injured groups. Our previous study also included quantification of other variables such as bone formation rate, osteoclast number and activity, voluntary activity, and levels of interleukin 6 (IL-6) in serum. These parameters were not measured in the current study, but we anticipate that we would observe similar trends in these outcomes following MI in mice.

The SNS acts through signaling via three types of β-adrenergic receptors (β_1_, β_2_, and β_3_). Activation of the SNS via adrenergic neurotransmitters generally inhibits osteoblast proliferation [16] and triggers osteoclastic bone resorption [17]. Several studies showed that treatment with non-specific SNS inhibitors increased bone mass [18, 19], and stimulation of the SNS decreased bone mass [20]. While all three β-adrenergic receptors are present in the skeletal system, β_2_ is prevalently expressed in both osteoblasts and osteoclasts [18, 21-28]. The specific role β_2_ plays with these two cell types have not been thoroughly investigated, but there is evidence that β_2_ stimulation has a significant deleterious effect on bone [18]. In contrast, relatively little is known about the role of β_3_ receptors in the skeletal system. β_3_ has been shown to increase osteoclastogenesis and subsequent bone resorption *in vitro*, but there is no consensus of its effect *in vivo* [21, 29]. Beta-blockers have been investigated as a potential treatment for the skeletal system in several studies. Daily treatment with a β_1_/β_2_ agonist triggered an osteoclastic response [18, 30, 31] with increases in RANKL and IL-6 expression; another study determined that treatment with a general beta-blocker lead to a high bone mass phenotype [32]. In the current study we utilized the same β_3_ adrenergic receptor antagonist (SR 59230A) that was able to lower protease activity, myeloid cell content, and mRNA levels of inflammatory cytokines in atherosclerotic plaques following MI in mice [10], thus allowing us to determine if the same underlying mechanisms contributed to bone loss following MI.

Our findings using β_3_-AR antagonist treatment in mice following MI yielded somewhat inconsistent results. In the study by Dutta et al. [10], treatment with the same β_3_-AR antagonist decreased the inflammatory response through increased withdrawal of stem cell retention factors by β_3_ expressing cells. Four days post-MI, blood HSPC’s levels and inflammatory markers were lower in treated groups [10]. In contrast, we observed greater monocyte and neutrophil levels in the blood of treated mice, although this could be largely due to the time point when blood was collected (day 4 in Dutta et al. vs. day 10 in the current study). Additionally, our results suggest that β_3_-AR antagonist treatment generally led to decreased bone volume and diminished systemic bone loss after MI; these data are consistent with a previous study investigating β-adrenergic blockade in rats during hindlimb unloading [33]. In this study, hindlimb unloading resulted in a 20% decrease in cancellous vBMD, but this bone loss was halved in rats treated with a general β-blocker (propranolol) through stimulation of osteoblastic activity and suppression of osteoclastic activity. This study similarly showed that β-blocker treatment of cage activity control rats resulted in net trabecular bone loss at the proximal tibia relative to vehicle-treated controls. We also observed effects of β_3_ inhibition on changes in lean tissue area and adipose tissue area. While it was expected that MI may decrease lean tissue and adipose tissue mass due to post-operation recovery, treatment with the β_3_-AR antagonist exacerbated the loss of both lean and adipose tissue. These findings are contrary to previous studies that showed that β_3_ agonists promote weight loss, especially in obese mice and rats [34-37].

This study established, for the first time, a novel mechanistic relationship between acute cardiac injury and subsequent remodeling in bone. Despite some epidemiological evidence supporting a link between Fx and MI, this is the first study to show a causal relationship between these two seemingly unrelated events. Interestingly, the epidemiological evidence linking incidence of Fx and MI suggests this link is bidirectional, therefore Fx may also exacerbate atherosclerosis, leading to increased risk of subsequent MI. This is supported by a study by Chiang et al., which reported significantly higher risk of subsequent MI following hip fracture (hazard ratio = 1.29) [38]. Further studies are required to establish this relationship mechanistically.

The current study has some limitations that must be acknowledged. First, quantification of monocytosis was performed by drawing blood through a needle, which allows for the possibility of cell lysing and subsequent inaccuracy in results. Furthermore, our method of measuring monocytosis was a more general assessment of blood cell composition than in Dutta’s study, which looked at many markers of inflammation in more specific areas such as the bone marrow and the spleen, two important areas for monocyte proliferation. Secondly, the β-blocker we used was specific to β_3_ receptors, where its interaction with the skeletal system is not as well known as the other types of β-adrenergic receptors. As a result, it is possible the effects we see with treatment could be due to the minimal presence of β_3_ receptors in bone cells. It is possible that a general β-blocker would be more successful in preventing bone loss following MI. Third, many of our variables were measured only at one time point (10 days post-MI). While we have previously established that peak bone loss occurs between 7-14 days after fracture [9], it is possible that 10 days post-MI is not the optimal time point to assess bone loss following this type of injury, and we did not quantify recovery of bone at later time points. Additionally, the animals used for this study were young, male mice, while MI and Fx occur more commonly in the older population. We have also shown that bone remodeling after fracture differs between young and middle-aged mice [9]. However, we chose to study young male mice as it allows for minimal confounding factors from existing comorbidities and for their higher survival rate following MI. Finally, we did not directly quantify bone formation or bone resorption rates in these mice, therefore it is difficult to determine the biological mechanisms underlying the observed changes in bone mass and microstructure.

## Conclusions

This study is the first to establish a causal relationship between MI and bone loss at multiple skeletal sites, and suggests that the SNS may have a governing role in this adaptation. This injury-induced response may also be operative in human subjects after MI, and may be a potentially catastrophic co-morbidity in post-MI patients. Further delineating the relationship and mechanisms governing this crosstalk could inform future treatments aimed at preventing injuries and preserving skeletal health following ischemic injuries.

## Acknowledgements

We would like to thank Dr. Heike Wulff for her valuable input on this project. Research reported in this publication was supported by the National Institute of Arthritis and Musculoskeletal and Skin Diseases, part of the National Institutes of Health, under Award Number AR071459. The content is solely the responsibility of the authors. The funding bodies were not involved with design, collection, analysis, or interpretation of data; or in the writing of the manuscript. The authors have no conflicts of interest to disclose.

